# Cellular and synaptic organization of the Octopus vertical lobe

**DOI:** 10.1101/2025.01.29.635406

**Authors:** Flavie Bidel, Yaron Meirovitch, Fuming Yang, Jeff William Lichtman, Binyamin Hochner

**Affiliations:** Department of Neurobiology, Silberman Institute of Life Sciences, The Hebrew University, Jerusalem, Israel; Department of Molecular and Cellular Biology, Harvard University, Cambridge, MA, United States

## Abstract

Understanding memory formation and its influence on behavior is a central challenge in neuroscience. Associative learning networks, including the mushroom body in insects, the cerebellum in mammals, and the vertical lobe (VL) in cephalopods, typically exhibit a 3-layered architecture, characterized by divergence (fan-out) followed by convergence (fan-in), facilitating sparse sensory coding (Babadi and Sompolinsky, 2014; Lin et al., 2014; Litwin-Kumar et al., 2017; Turchetti-Maia et al., 2017). Previously, using volumetric electron microscopy, we showed that the VL uniquely comprises 22 million simple amacrine (SAM) interneurons, each receiving a singular input subject to activity-dependent long-term potentiation, contrasting with typical middle-layer interneurons (Bidel, Meirovitch et al., 2023). We also demonstrated that these SAMs provide excitatory feedforward input to the output cell layer, balanced by approximately 400,000 inhibitory complex amacrines (CAM), which are morphologically diverse and integrate numerous inputs (Bidel, Meirovitch et al., 2023). Here, we leverage the same digital tissue to explore the CAMs’ morphological diversity, identifying correlations between structure, postsynaptic site density, and synaptic input proportions, which led to the classification of CAMs into distinct groups. Further analysis of the input layer in the VL revealed a meticulous structural and synaptic compartmentalization, with distinct synaptic bouton types forming three zones that integrate different inputs towards CAMs. Additionally, we identify the potential presence of a neurogenic niche in the VL, hinting at parallels with neurogenic processes in other species and warranting further investigation, particularly in the context of learning and memory. This study deepens our understanding of the VL’s cellular and synaptic architecture, revealing both shared and unique features compared to other associative networks, and highlighting the intricate interplay of structural and functional elements in memory formation.

**Highlights (85 characters including space):**

- The VL both aligns with and deviates from standard associative network features
- The VL input layer presents a unique compartmentalization by axonal bouton type
- The minority inhibitory interneurons differentiate into unique functional subtypes
- Mirroring the mammalian hippocampus, the VL may contain an adult neurogenic niche

## Results and discussion

The VL is a critical center for associative learning in cephalopods (Boycott and Young, 1955; Sanders, 1975; Fiorito and Chichery, 1995; Shomrat et al., 2008, 2015; Turchetti-Maia et al., 2017). Our previous work mapped 516 cellular processes in the VL, revealing two interconnected feedforward networks (Bidel, Meirovitch et al., 2023). These networks are influenced by two primary afferents: the superior frontal lobe axons (SFL) and the complex input neurons (CIN), which predominantly target two distinct amacrine interneuron populations in the VL’s input layer. The simple amacrines (SAMs) and the complex amacrines (CAM) differ significantly in morphology and connectivity, with the SAMs characterized by a single input and the CAMs by their integration of multiple diverse inputs. This study further explores the VL network, utilizing previously acquired electron microscopy (EM) imaging data and segmentation data (available at https://lichtman.rc.fas.harvard.edu/octopus_connectomes) to further characterize cell types and uncover the intricacies of their synaptic and cellular organization.

### Morphological diversity of CAMs reflects subtypes specialized for a type of sensory inputs

Within the VL, two primary interneuron types are discernible: the 22 million excitatory SAMs, which are uniform in nature, and the 500,000 inhibitory CAMs. Although CAMs represent only about 2% of VL cells, they exhibit distinctive groups of morphological diversity (Figure 1A). This suggests that despite their scarcity, they play a pivotal role in the VL network function. SAMs have a simple morphology, possessing a single non-bifurcated dendritic trunk and a unique ‘palm’-like structure that likely receives a distinct visual feature from a single SFL neuron via a large presynaptic bouton. In contrast, CAMs display expansive dendritic arborizations and a clear separation between input and output regions, demonstrating substantial anatomical diversification. To identify possible functional classes of CAMs, we classifiedthe 53 reconstructed CAMs based on their gross morphological features, postsynaptic dendritic branching, and input distribution patterns (see Methods). Our analysis revealed two primary dendritic branching orientations: horizontal CAMs, which extend their dendritic processes laterally within the SFL tract, and vertical CAMs, characterized by a primary inward-directed neuritic backbone (Figure 1A). For the horizontal CAMs (Figure 1A-right panels), we identified two subtypes: The “flag” (Fl) cells (n=8) are characterized by a spatially compact and densely branched arborization. In contrast, the “long-range” (Lr) cells (n=4) exhibit a sparser arborization, indicating fewer branches and a less compact structure. The vertical CAMs (Figure 1A-left panels), can be further categorized into three distinct types: The “thorny cells” (Th) (n=3) are characterized by numerous tiny membranous protrusions, reminiscent of vertebrate dendritic spines, that emanate from their primary neuritic backbone and second-order branches. The “very narrow dendritic cells” (Vw) (n=11) predominantly exhibit the less extensive dendritic fields among all CAMs (4.64 +/- 2.64 μm, extension from the main neuritic) and have sparsely branched neuritic backbones. Some even display dual backbones originating from their cell body. Lastly, the “narrow dendritic cells” (Nw) (n=9) feature a slightly more expansive dendritic field that is denser than that of the very narrow dendritic cells.

**Figure 1:**
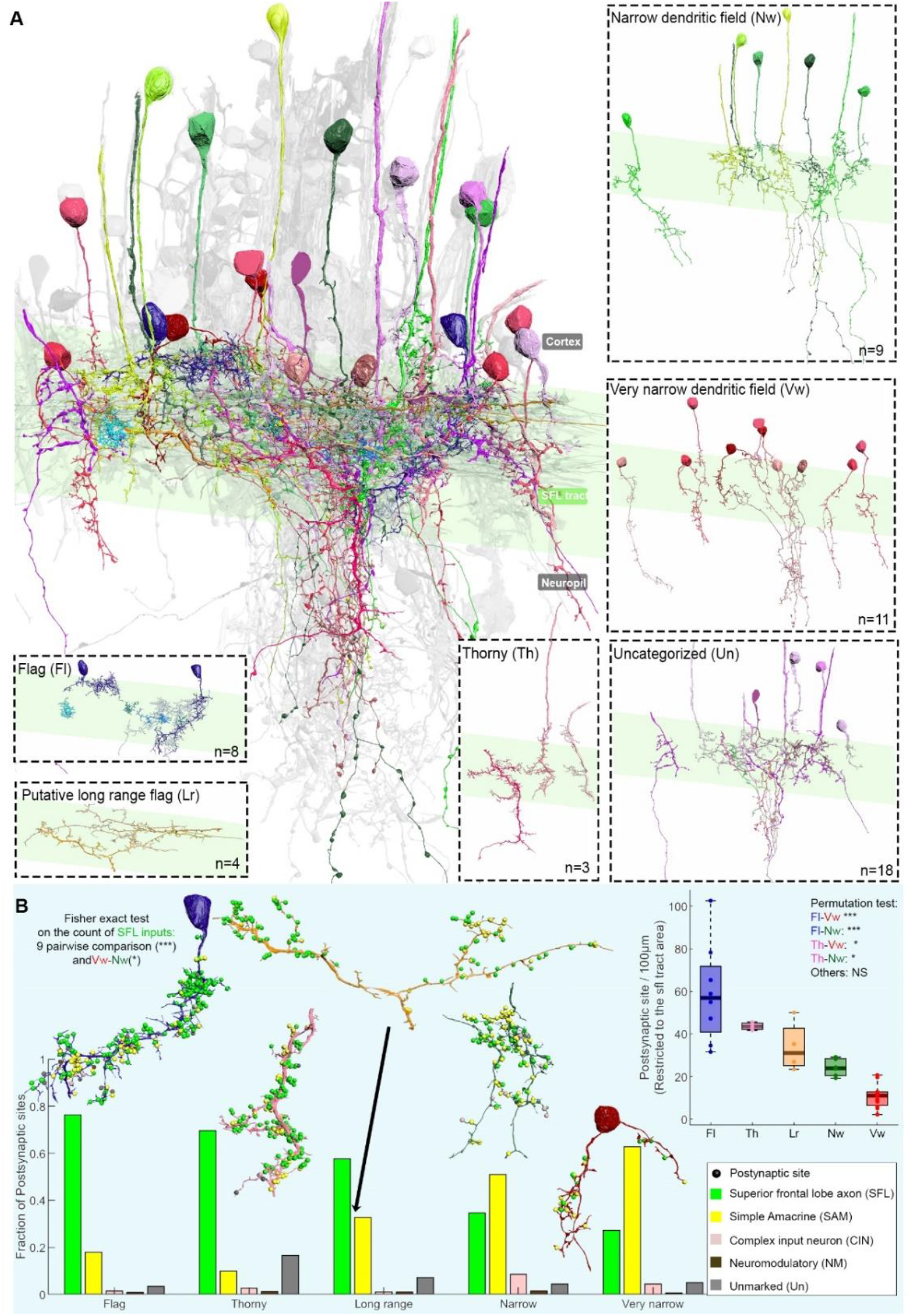
The CAM can be roughly classified into 5 morphological subtypes. A. A 3D reconstruction of the 53 CAMs, color-coded by subtype, is overlaid on a grayscale representation of the 516 reconstructed cells, illustrating their extensive ramification within the SFL tract area (highlighted in green). CAMs display a broad spectrum of morphological subtypes, bifurcating into two primary classes: horizontal cells, encompassing the flag (Fl, in blue) and putative long-range (Lr, in orange) cells, and vertical cells, which include the narrow dendritic (Nw, in red), very narrow dendritic (Vw, in green), and thorny (Th, in pink) cells. Each subtype is spotlighted within a dashed square panel. Notably, several cells remain unclassified (Un, in gray) due to their limited presence within the volume or insufficient observations for categorization. B. Morphological differences among CAMs correlate with variations in synaptic input proportions, with the analysis specifically restricted to inputs within the SFL tract for normalization purposes. All Fisher exact tests with Bonferroni correction, based on the count of SFL inputs per subtype, yielded significant results. A 3D model for each subtype is presented, with postsynaptic input sites (puncta) color-coded based on their presynaptic partner. The postsynaptic site density within the SFL tract area varies significantly among subtypes (Kruskal-Wallis test, p<0.001). A permutation test indicated that flag and thorny cells possess notably higher synaptic densities compared to narrow and very narrow dendritic field cells. Levels of statistical significance are denoted by asterisks: *** p<0.0001, ** p<0.001, * p<0.005.

Note that given the pronounced difference in diameters between the neuritic backbones (208.27 +/- 87.95 nm) and axons (121.55 +/- 21.28 nm), it’s probable that CAMs output is propagated actively though long thin spiking axonal processes. As common in invertebrate neurons, the nonlinear “integrate and fire” zone, take place at the spike initiation zone at the transition between the dendritic and axonal segments (Bullock and Budelmann, 1991).

While volume EM is invaluable, it poses challenges for extensive cell type classification based on morphometric measurements due to the truncation of cells within our volume. However, we supplemented our classification with statistical analysis of neuritic backbone diameters, a crucial parameter influencing electrical properties like the electrical length constant. This analysis revealed significant differences in backbone diameters across CAM subtypes, with thorny cells having the largest (331.55 ± 159.31 nm, n=16 from 3 cells) and flag and very narrow CAMs the smallest (165.89 ± 53.56 nm and 153.04 ± 92.22 nm, respectively). This suggests distinct electrical properties; indeed, cell types that seem to integrate inputs from over large distances have thicker main dendritic diameters, likely contributing to a longer electrical space constant required for the integration of widely spread postsynaptic inputs (Figure S1).

Flag CAMs (n=8) and narrow dendritic subtypes (n=9+11) are the two subtypes that display the most significant morphological differences. Flag cells have dense, bushy dendritic fields that cover a condensed volume within the SFL axonal tract (Figure 1A). In contrast, the backbone neurites of the narrow and very narrow CAMs cross the SFL vertically into the deeper neuropile, emitting sparse and relatively short secondary branches. This group overlaps a narrow and elongated volume of the SFL tract and continues to receive SAM inputs in the deeper neuropile below the tract. Narrow CAMs are thus tuned to integrate relatively more inputs from SAMs than SFL axons.

These morphological differences are indeed reflected in the synaptic input patterns of CAM subtypes within the SFL area. When focusing on these patterns, there are marked differences in both input density and the proportions of these inputs (p< 0.05, Figure 1B). For instance, flag CAMs register an average of 60 postsynaptic sites per 100 μm within the SFL-VL tract, comprising 79% of SFL and 19% of SAM inputs. Conversely, very narrow CAMs display a lower synaptic density, with 10 postsynaptic sites per 100 μm, predominantly influenced by SAMs (63%) and SFL (27%) (Figure 1B).

In addition to the excitatory glutamatergic SFL axons and cholinergic SAM inputs described above, all CAMs consistently receive inputs from two additional distinct sources: Complex Input Neurons (CIN), which exclusively innervating CAMs and are suggested to convey inhibitory signals into the CAMs, and a small population of neuromodulatory inputs (Bidel et al., 2023). While the proportion of SFL and SAM excitatory inputs varies significantly among CAM subtypes, the proportion of CIN inputs shows less variation, suggesting that inhibitory input is tuned in all CAM subtypes to balance the total excitatory inputs.

In summary, the morphological diversity of CAM subtypes, including the density, volume, and orientation of their dendritic fields, determines their connectivity patterns and the type of information each subtype integrates. The broad horizontal reach of flag CAMs intersects with numerous SFL axons and a varied set of SAMs. On the other hand, vertically oriented CAMs, like the very narrow CAMs, likely connect with a limited set of SFL axons and a specific SAM group. These structural distinctions not only lead to varied integration of visual inputs and input densities among CAM subtypes but also hint at their functional specializations.

As CAMs were proposed to monitor the VL’s global input activity (Bidel et al., 2023), our results suggest that different CAM subtypes are tailored to integrate different patterns of global activities, which are represented by different assemblages of SFL axons and groups of SAMs. The main connectivity difference between CAM subtypes lies in the dichotomic differences in the SFL inputs relative to those of the SAMs (Figure 1B). It is feasible that each CAM subtype monitors different combinations of ongoing activity, possibly implemented in different local microcircuits (submodules). This may be important for processing stimulus-specific information stored at the monosynaptic SFL-to-SAM connections. For instance, CAMs with larger spans, such as long-range, flag, and thorny, might be involved in diverse assemblies of SFL and SAMs. In contrast, CAMs with shorter spans, such as narrow and very narrow, appear to be dedicated to specific, spatially compact association modules (Bidel et al., 2023).

While our current categorization provides foundational insights, the need for broader reconstructions and advanced techniques. Future studies, employing cutting-edge algorithms and comprehensive reconstructions combined with molecular atlases and in situ hybridization, will be pivotal in decoding VL cell diversity and function.

### The VL input layer is organized in three clusters of distinct synaptic glomeruli

The CAMs predominantly receive their inputs within the SFL tract area that form the VL input layer. These inputs are supplied by axonal boutons from both the SFL and CIN, as well as by the specialized “palm” and trunk swellings of the SAMs. Intriguingly, the SFL inputs are unique in that they come from two functionally distinct types of SFL boutons: the small SFL boutons, which target exclusively the CAMs, and the large SFL boutons, which not only connect to one SAM palm but also frequently form triadic arrangements with the innervated CAM (Bidel et al., 2023).

Upon inspecting the electron micrographs, the clustering of boutons and palm enlargements within the SFL tract area is immediately evident. This region showcases a profusion of varicosities brimming with densely packed vesicles. Notably, varicosities of similar types tend to group together (Figure 2A). To gain a deeper understanding of the spatial arrangement within these synaptic zones, we reconstructed all neighboring boutons over an area spanning approximately 15,000 μm^3^ in the SFL tract. Our findings underscored the intricate organization of these varicosity-dense synaptic zones. We discerned that the two SFL bouton types—small and large —each form spatially distinct spatial clusters (Figure 2B, C). Additionally, neuromodulatory boutons (NM), the CIN axonal boutons, and other yet-to-be-characterized boutons collectively constitute a third distinct cluster (Figure 2C To further investigate the structural compartmentalization of the VL input layer, we selected representative areas from each of these clusters and performed saturated cubic reconstructions (125 μm^3^) for each (Figure 2E). These cubes, labeled 1-3 in Figure 2E, are representative samples from the aforementioned clusters, allowing us to delve into the specific connectivity differences within each cluster. Each region revealed its own unique architectural layout. showcasing distinct variations in both neuronal composition (Figure 2F) and connectivity patterns (Figure 2G, H, I).

**Figure 2:**
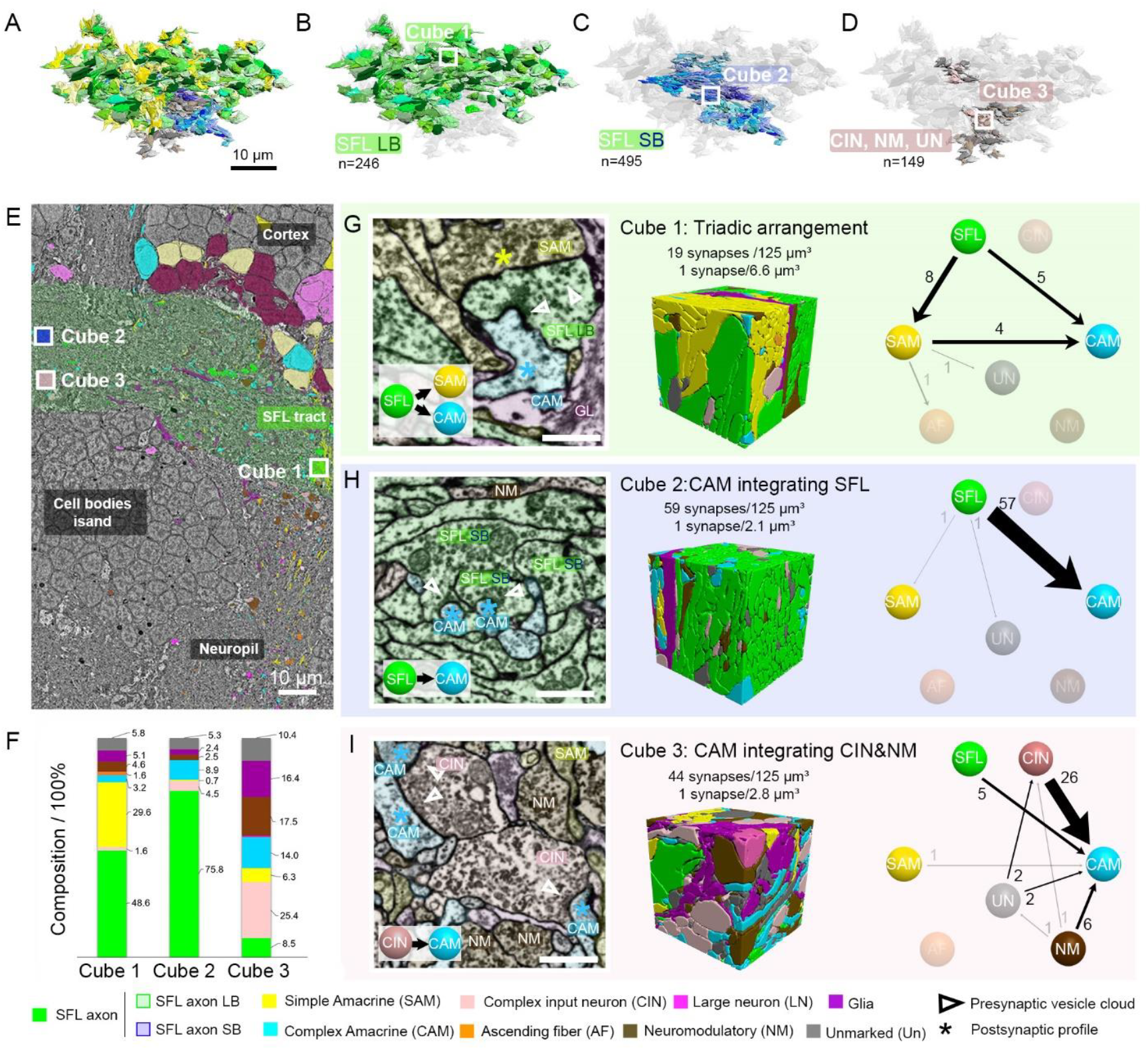
Topological organization of the VL input layer demonstrates distinct synaptic compartments formed by clustering varicosities of similar nature. A. 3D reconstruction of adjacent varicosities, color-coded by type, within an approximate 15,000 μm^3^ volume of the SFL tract, revealing three distinct clusters. B. 246 large SFL boutons depicted in varying shades of green. C. 495 small SFL boutons illustrated in different shades of blue. D. A combined representation of 149 CIN (in pink), NM (in brown), and uncharacterized boutons (labeled as UN). E. Low-resolution electron micrograph (EM) of the VL section used for the reconstruction. Highlighted squares (labeled Cube 1-3) pinpoint the locations of the three 125μm^3^ cubes subjected to saturated reconstruction and synaptic identification. F. Volumetric analysis unveils diverse compositions based on the proportion of neuronal elements present. G, H, and I. Each panel showcases an EM highlighting the predominant synapses of the respective cube (Scale bar = 500nm), a 3D reconstruction of the cube color-coded by cell type, and the connectivity diagram within that cube. Light gray and transparent elements signify minor connections and less prevalent neuronal elements within the cube. Collectively, the three cubes elucidate three unique patterns of synaptic connection areas.

We subsequently delved into the structural compartmentalization of the VL input layer by executing a saturated cubic reconstruction (125 μm^3^) across these three distinct areas (Figure 2E).

The first cube (Figure 2G), which denote as the “Triadic arrangement” zone, displayed the prominent presence of SFL LB and SAM palms. The volume of this cube was predominantly composed of SFL processes (50%) and SAM processes (30%). Out of the 19 synapses identified within the cube, 8 were between SFL and SAM. In contrast, synapses from SFL-to-CAM (5/19) and SAM-to-CAM (4/19) were less frequent. These three types, namely SFL, CAM, and SAM, were locally interconnected in a triadic configuration. Given the extensive nature of the compartments associated with these synapses and considering the large size of the SFL large bouton-SAM palm synapse, the observed synaptic density in this region was relatively low, with an average of 1 synapse per 6.6 μm^3^. The substantial volume of these compartments, estimated to contain ∼2500-5000 vesicles in the SFL presynaptic area, suggests a high reliability in synaptic transmission within this zone.

In the second cube (Figure 2H), which represents the CAM integrating SFL zone, SFL axons, especially the small boutons, occupied a significant 76% of the cube’s volume. Despite CAM processes constituting only about 9% of the cube’s volume, they were the primary target of these SFL axons, receiving 57 of the 59 synapses identified within the cube. This specific region, where CAMs are the focus, had a synaptic density of 1 synapse per 2.1 μm^3^. This density is notably half as dense as what has been observed in the neocortex (Kasthuri et al., 2015).

For the third cube (Figure 2I), which is indicative of the CAM integrating CIN and NM area, the combined volume of Nm, CIN, and unidentified processes made up 53% of the cube’s space. CAMs and SFL processes took up 14% and 8% of the volume, respectively. Within this cube, all the identified synapses were oriented towards the CAM, with a synaptic density of 1 synapse per 2.8 μm^3^. Breaking down the synaptic relationships with the CAM, we found CIN-to-CAM connections accounting for 26 out of the 44 synapses, Nm-to-CAM for 6 out of 44, and SFL-to-CAM for 5 out of 44.

Additionally, we observed that astrocyte-like glial processes seemingly encapsulate these distinct zones (refer to glial processes, purple, in cubes 1, 2, and 3; Figure 2G, H, I). This encapsulation might offer functional synaptic isolation between zones with differing roles. For instance, such glial demarcation could potentially prevent retrograde messengers like NO (Turchetti-Maia et al., 2018), known to play a role in LTP within the VL tract, from migrating between different synaptic regions.

In summary, our investigations into the SFL tract area have unveiled a pronounced structural and synaptic compartmentalization, demarcating it into three distinct regions. Each of these regions is characterized by a unique spatial differentiation of inputs directed to the CAMs, suggesting specialized roles in information processing. The connectivity motif of the first region, the only one that includes SAMs (Figure 2G), appears to be overarching of the network input activity. It potentially monitors the stimulus-specific and nonspecific activities at the input layer of the association module including its modifications due to activity-dependent short- and long-term gain control modifications. The second region (Figure 2H), in contrast, seems poised to monitor the background SFLs input. Intriguingly, if the small SFL boutons lack long-term potentiation the input-output relationship at this zone might remain independent of activity-dependent modulations, offering a genuine reflection of the ongoing SFL input activity. The third region (Figure 2I) stands out in its possible role of integrating external supervising inputs of inhibitory (pain) and neuromodulatory (supervising) signals(Bidel et al., 2023; Turchetti-Maia et al., 2017). Collectively, these findings underscore the CAM’s role in continuously overseeing the VL’s activity, with each region dedicated to processing specific types of spatially organized information. Collectively, these findings underscore the CAM’s role in setting microcircuit motifs, with each region monitoring different computationally relevant activities within the VL.

### Evidence for the existence of an adult neurogenic niche in the Vertical lobe

The VL cortex predominantly consists of three intrinsic neuron types, collectively making up about 90% of its cellular composition: the ubiquitous SAMs (89.3%), the morphologically diverse CAMs (1.6%), and the less common large neurons (0.68%). Intriguingly, an additional 8.42% of the VL cortex’s cells deviate from these established neuronal profiles. These cells display unique morphological and ultrastructural characteristics and notably lack clear synaptic connections (Bidel et al., 2023). While some align with recognized glial types, such as radial glia and astrocyte-like glia, a significant portion in the VL’s inner cortex remains unidentified. Notably, within this unidentified segment, there exists a distinct cluster of cells that congregate within the inner cortex of the VL (Figure 3A) that we explored here.

**Figure 3:**
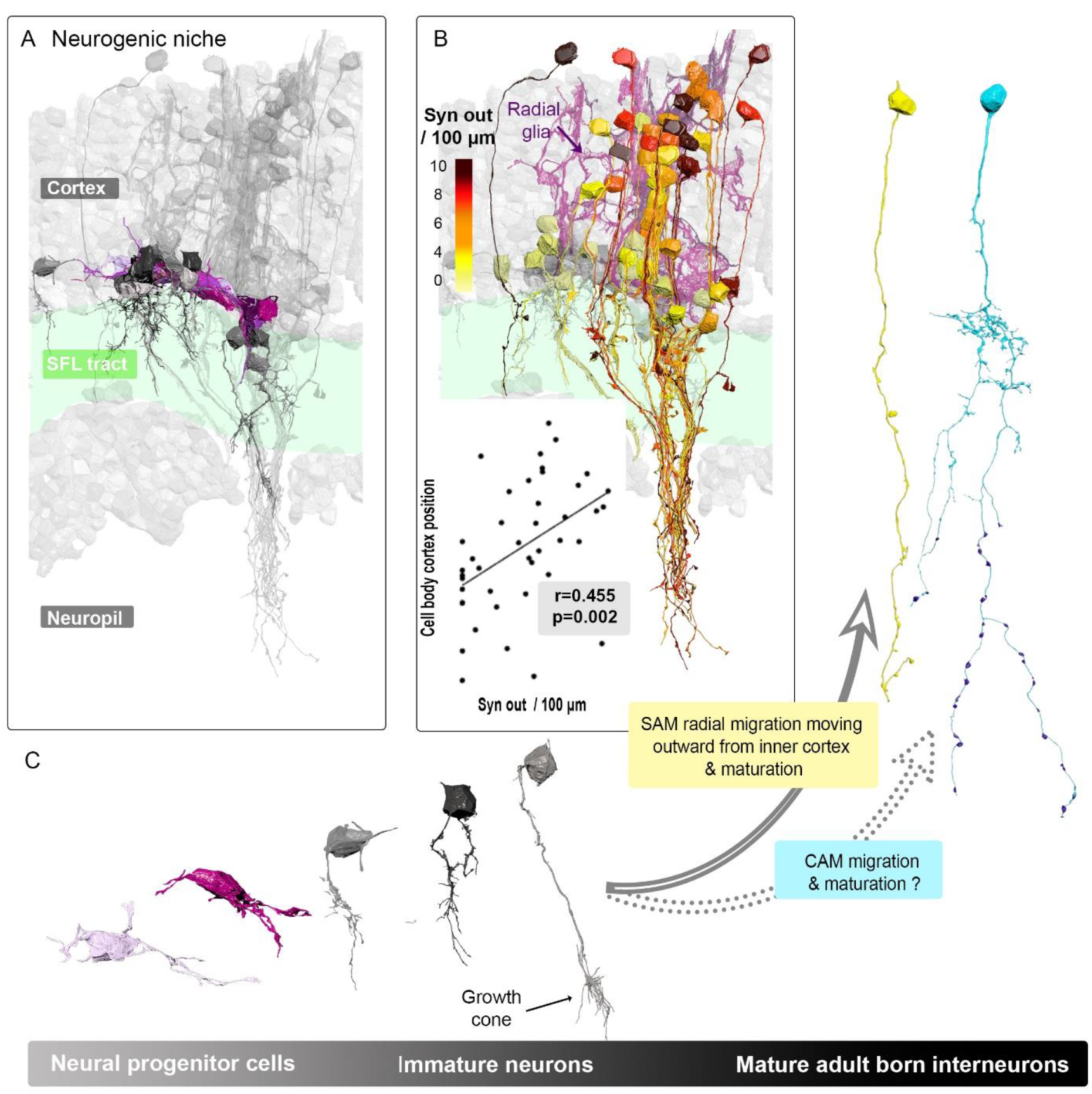
Adult Neurogenic Niche in the VL: Cellular Composition and Organization. A. Reconstructions of putative neurogenic cells (n=24) in the inner cortex, highlighting neural progenitor cells (purple) and immature neurons (gray), representing a continuum of differentiation. B. SAMs reconstructions (n=43) color-coded by synaptic output density (syn out per 100um normalized within the SFL tract) positioned above the neurogenic niche and adjacent to radial glia. The bottom-left plot shows the significant Pearson correlation between cell body position and synaptic output density, indicating SAM maturation. C. Schematic representation of the VL’s adult neurogenic niche organization and composition.

Our 3D reconstructions revealed two distinct cell types that diverge from the characteristics of mature neurons. The first, which we’ve termed “Neural progenitor cells”, possess elongated cell bodies with often irregular and folded nuclear membranes. These cells also exhibit filopodia and membranous lamellipodia-like extensions, seemingly wrapping around similar cells. Their absence of synaptic connections, combined with their morphological and ultrastructural resemblance to early postnatal developing granule cells in the cerebellar cortex, suggests they might be progenitors of amacrine cells (Cervantes et al., 2022).

Positioned above these progenitor cells, we identified another set of cells with more conventional spherical neuronal morphologies. These cells, potentially immature neurons, extend trunks into the neuropil, frequently devoid of synaptic connections. These trunks are enriched with grayish filaments, possibly filamentous actin, hinting at sites of plastic reorganization. Some even culminate in growth cone-like filopodia, further suggesting their developmental nature. Together, these Neural Progenitor cells and immature neurons seem to form an adult neurogenic niche within the VL (Figure 3A). This niche mirrors mammalian neurogenic regions found in the dentate gyrus of the hippocampus and the subventricular zone (review in Bartkowska et al., 2022).

To bolster this hypothesis, we assessed the synaptic output density of SAMs based on their cortical position, using it as an indicator of neuronal maturity. Our analysis unveiled a significant correlation between synaptic output density and cortical position (r^2^= 0.455, p=0.002), suggesting a potential neuron maturation along a radial path, moving outward from the inner cortex and aligns with the presence of radial glial fibers in the VL cortex (Figure 3B).

Our study has uncovered evidence suggesting ongoing integration of new interneurons, predominantly SAMs and potentially CAMs, into the VL of adult octopuses (Figure 3C). This observation, which may indicate neurogenesis in the VL, raises intriguing questions about the role of these newly integrated neurons in the octopus’s nervous system, particularly given the rapid growth associated with the short lifespan of *Octopus vulgaris* and concomitant fast and large increase in VL volume and cell number (Budelmann, 1995; Young, 1963). Yet it is still tempting to draw parallels with mammalian systems, where neurogenesis in structures like the hippocampus and olfactory bulb is linked to learning and memory processes (Gould et al., 1999; Rochefort et al., 2002; Sakamoto et al., 2011; Shani-Narkiss et al., 2020; Vicidomini et al., 2020), the possible role of neurogenesis in the VL learning memory remains speculative and require further studies.

In the context of the VL, it is conceivable that the addition of new neurons could contribute to the dynamic nature of the octopus’s nervous system, accommodating its rapid physical growth. However, the direct involvement of these new neurons in learning and memory within the VL is less clear. Previous studies have indicated that while long-term potentiation (LTP) in the VL is crucial for learning (Shomrat et al., 2008; Turchetti-Maia et al., 2017), it may not be directly tied to long-term memory storage, which is believed to occur outside the VL(Boycott and Young, 1955; Shomrat et al., 2008). Additionally, the unique characteristics of LTP in the octopus VL, such as its reliance on a NO-dependent mechanism, independent of de novo protein synthesis persistent activation of (Turchetti-Maia et al., 2018). This mechanism further complicate relating the VL LTP with later phases of long-term memory acquisition and the comparison with mammalian systems.

The driving goal of this work was to provide a detailed view of the cellular and synaptic organization of the octopus vertical lobe to date. We therefore schematized a spatial map of the vertical lobe (Figure 4) to summarize this organization and illustrate its different features. This map shows the intricate interplay between various neuronal elements, highlighting the distinct regions and their respective functions. From the densely packed SAMs to the morphologically diverse CAMs, and the potential neurogenic niches, each component plays a pivotal role in the VL’s complex information processing. The topological organization of the VL input layer, with its synaptic compartments and glial-defined boundaries, underscores the VL’s unique evolutionary trajectory. Furthermore, the presence of both shared and distinct features compared to classical associative learning networks emphasizes the VL’s significance in understanding neural evolution and function. This comprehensive spatial representation serves as a foundational reference for future studies.

**Figure 4:**
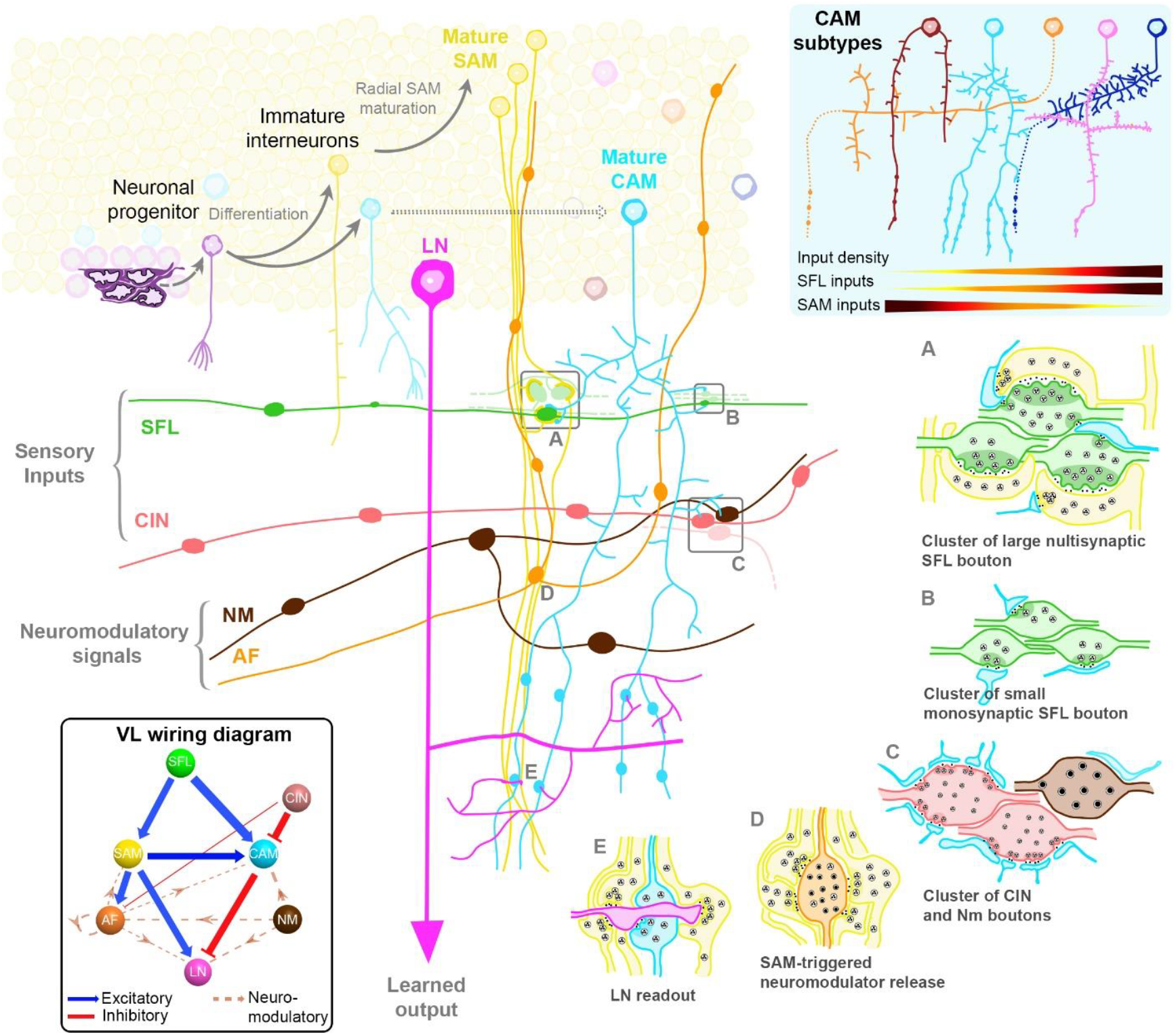
Synaptic and Cellular Blueprint of the Vertical Lobe. This schematic captures the intricate organization of the Vertical Lobe (VL) along with its wiring diagram (modified from Bidel, Meirovitch et al. 2023). Two primary afferents, the Superior Frontal Lobe Axons (SFL) and the Complex Input Neurons (CIN), convey secondary visual cues and potential “pain” signals, respectively. They innervate two distinct amacrine interneuron populations within the VL’s input layer, the SFL tract. This layer shows a meticulous organization, with like boutons clustering together, forming distinct synaptic glomeruli clusters. The Simple Amacrines (SAMs), constituting the majority at around 23 million, stand out with their singular input and simplicity in morphology and connectivity. In contrast, the Complex Amacrines (CAMs) represent only 400 000 cells of the cortex, and exhibit a rich morphological diversity, with various subtypes each having unique input densities and fractions. A potential neurogenic niche in the inner cortex suggests the capability for continuous integration of new amacrines throughout the octopus’s lifespan. These interneurons converge onto the Large Neuron Processes (LN), the sole VL output, within the neuropile. The circuit is further enhanced by a vast modulatory network, including the Ascending Fibers (AF) and the Neuromodulatory Fibers (NM).

## Supplementary figure

**Figure S1:**
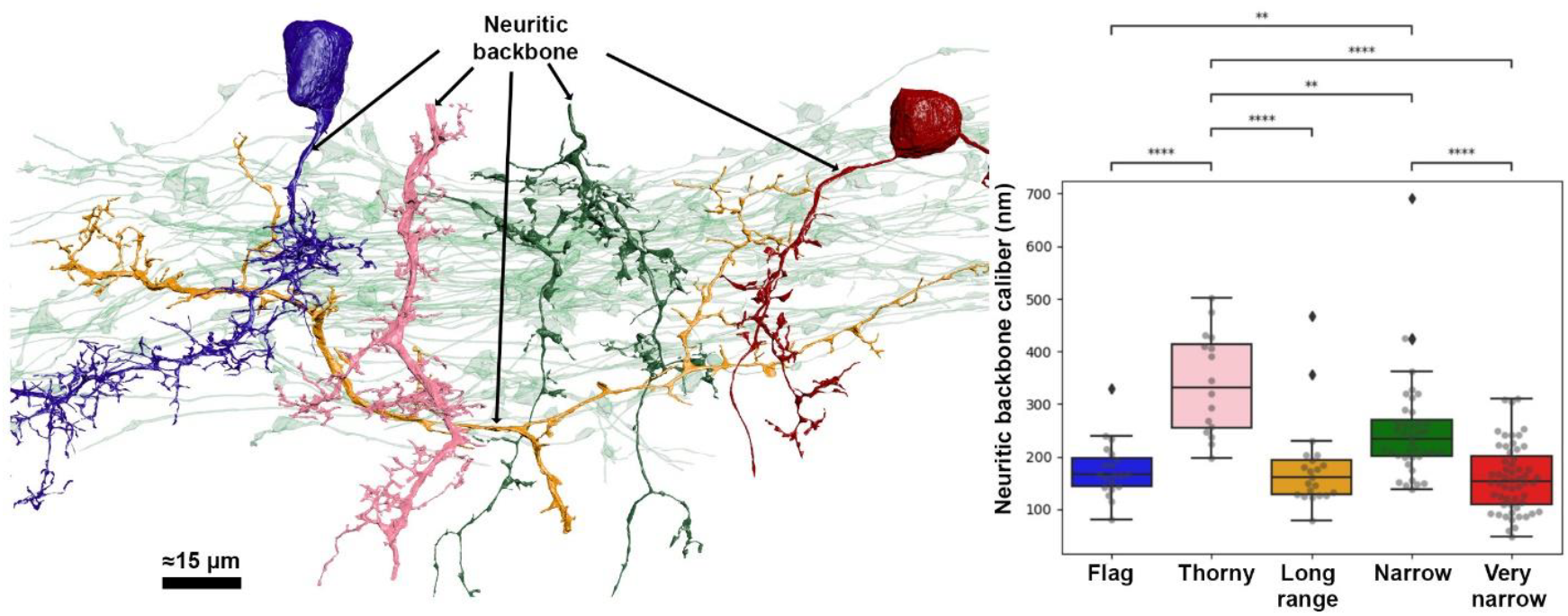
The 5 morphological subtypes of CAMs display different backbone diameters, suggesting variations in their electrical properties. A. Reconstructions of CAMs, color-coded by subtype, are overlaid on the SFL tract (green). CAMs exhibit a wide array of morphological subtypes. These bifurcate into two primary classes: horizontal cells, which include the flag (Fl, in blue) and putative long-range (Lr, in orange) cells, and vertical cells, which comprise the narrow dendritic (Nw, in red), very narrow dendritic (Vw, in green), and thorny (Th, in pink) cells. The neuritic backbone significantly varies among subtypes (Kruskal-Wallis test, p<0.001). A permutation test with Bonferroni correction indicated pairwise differences. Levels of statistical significance are denoted by asterisks: *** p<0.0001, ** p<0.001, * p<0.005.

## Method

### Method Details

The acquisition of the VL EM image volume and segmentation method employed in this study were previously detailed in Bidel, Meirovitch et al., 2023. In brief, an adult Octopus was anesthetized in seawater containing 55 mM MgCl2/2% ethanol and then fixed in 3% glutaraldehyde, 0.065 M phosphate buffer, 0.5% tannic acid, and 6% sucrose for 12 hours at 4°C. Following fixation, the samples were sliced into 1 mm sections using a vibratome and stained using a modified en bloc staining protocol. This entailed sequential staining with 2% osmium tetroxide, 0.5% thiocarbyhydrazide, 2% osmium tetroxide once again, and 4% uranyl acetate. Subsequently, the vibratome sections were dehydrated with acetonitrile, embedded in EPON, and sliced into 30 nm sections using an automated tape-collecting ultramicrotome. A total of 892 sections were mounted on 100 mm silicon wafers and imaged using an FEI Magellan Scanning Electron Microscope equipped with custom image acquisition software (Hayworth et al., 2014). The acquired images had a pixel size of 4 nm. The ‘connectome dataset’ is publicly available using (browser-based) Neuroglancer at https://lichtman.rc.fas.harvard.edu/octopus_connectomes with 8 nm resolution along with the connectomic segmentation.

For image annotation, including cell tracing and synapse labeling, we employed VAST (Berger et al., 2018), utilizing a combination of manual annotation and machine learning-assisted segmentation (Pavarino et al., 2023). To generate 3D renderings of the traced objects, we created 3D surface meshes of segmented objects using VastTools (developed in MATLAB) and imported them into 3D Studio Max (Autodesk Inc.).

### Criteria for Cell Classification and synapse identification

In line with our previous research, we applied specific criteria for cell classification (Bidel et al., 2023). Afferent axons were differentiated by characteristics such as a constant radius, exclusive presynaptic boutons housing small synaptic vesicles, and no connections to cell bodies. Two subtypes of afferent axons, SFL axons, and CIN, were classified according to established criteria. VL interneurons were categorized based on their linkage to the SFL and their recognized intrinsic presence within the lobes. Within this category, we distinguished two types of AM cells, SAM and CAM, each serving distinct roles within the network. Efferent fibers primarily constituted LNs, characterized by cell bodies situated in the cortex and radial projections into the neuropil, often forming dendritic collaterals within the SFL tract. Neuromodulatory processes were recognized by the presence of large vesicles. Two main types of neuromodulatory processes, ascending fibers (AF) and neuromodulatory fibers (NMs), exhibited unique features. The glial category encompassed radial glia, astrocyte-like glia, and fibrous glia, each identified by specific structural attributes and their absence of synaptic connections. Lastly, cells categorized as “others” included elongated and spherical cell bodies, resembling progenitor-like cells, without evidence of cell division. For a more comprehensive understanding, readers are referred to Bidel et al. (2023) for detailed criteria.

Synapses were identified based on invertebrate synaptic characteristics, such as a vesicle cluster near the presynaptic membrane. We categorized them as traditional chemical synapses with small vesicles (fast transmission) or modulatory synapses with large vesicles. No electrical connections or clear postsynaptic densities were observed. All synapses lack clear postsynaptic density (PSD).

### Segmentation and Reconstruction

To investigate the morphological diversity of CAMs and track the maturation of SAMs along the radial cortex axis, we employed the same segmentation and annotation files utilized in our previous study.

For an in-depth examination of spatial organization within these synaptic zones, we meticulously reconstructed all adjacent boutons spanning an approximate volume of 15,000 μm^3^ within the SFL tract. Each bouton underwent categorization based on ultrastructural features, including vesicle size, mitochondria characteristics, and synaptic connections. SFL boutons were characterized by synaptic vesicles measuring 50-60 nm in diameter, accompanied by numerous small, round mitochondria. Among SFL boutons, two distinct types emerged: the SFL large boutons, consistently engaged in synapses with at least one SAM palm, and the small boutons, exclusively forming connections with single CAMs. Conversely, CIN boutons displayed the smallest vesicles within the VL, approximately 30-40 nm in diameter. They exhibited diverse mitochondria shapes, often of considerable size, and exclusively established synapses with the clear postsynaptic profiles of CAMs. Neuromodulatory boutons, encompassing NM and AF, featured larger vesicles, occasionally containing dense cores, and were larger than afferent axon boutons. However, they infrequently established synaptic connections. Boutons that couldn’t be definitively attributed to any of these categories were classified as “other.”

To assess tissue organization and synaptic density in various regions of the vertical lobe, we conducted four cubic reconstructions: three within the SFL tract area and one in the neuropil. The selection of each cube was guided by hypotheses formed from distinct observations of these regions. Within each 5×5×5 μm (125 μm^3^) cube, we expeditiously segmented all individual cellular processes using the collecting tool layer. A single annotator assigned cell identities, including SFL, CIN, SAM, CAM, AF, NM, LN, GLIA, or Unmarked. Additionally, all synapses underwent manual annotation, specifying their pre and postsynaptic partners, facilitating the quantification of synapses per type of connection. Connectivity diagrams within each cube were generated using Adobe Photoshop 2020. Each segmented cube was exported for reconstruction and volume measurements using the VastTools package.

### CAM Process Caliber and Distance Estimation

We employed a multi-step approach to estimate the caliber of CAM neuritic backbone processes. For this study, the neuritic backbone was defined as the main trunk connected to the cell body in most subtypes. However, due to the truncation of cells in the volume, the cell body was not always captured in the EM data for certain subtypes, such as CAM ‘Long Range’ and ‘Thorny’. In these cases, we approximated the neuritic backbone based on its appearance.For each CAM subtype, we used 4 to 11 cells, depending on the number of cells present in the volume that had sufficient length and were clearly identifiable by the annotator. In some cases, the limited number of cells available for certain subtypes restricted the sample size. For each of these cells, we randomly selected 3 to 5 measurement points along the neuritic backbone to best represent the overall diameter along its length. In total, measurements were taken from 30 cells across 5 different CAM subtypes, resulting in 153 diameter measurements.

Each location where the caliber was to be estimated was represented by a node in VAST, straddling the neuronal process. To determine the caliber precisely, we adjusted the coordinate to the medial axis of the process, achieved by performing a distance transform. This method calculates the distance from each object pixel to the surrounding tissue, identifying local maxima within a radius of 1.5 μm from the initial position. The process caliber was then defined as the distance from this medial axis to the nearest pixel outside the neuronal process

## Data analysis

All data were analyzed and visualized using MATLAB. Prior to statistical analysis, we tested the data for normality and homoscedasticity. Due to the non-normality of the data, we exclusively employed non-parametric statistical tests. For the comparison of CAMs SFL synaptic input and CAM backbone caliber, a non-parametric Kruskal-Wallis test was used. Upon rejection of the null hypothesis (p < 0.05), post-hoc exact permutation tests for independent samples were conducted. The resulting p-values were adjusted using the sequential Bonferroni correction method, as outlined by Holm (Holm, 1979). For the investigation of SAMs and the potential correlation between the synaptic output density of SAMs and their cortical position, a Pearson correlation analysis was performed. Throughout the manuscript, various measurements are presented as median +/- interquartile range due to the non-parametric nature of our data.

## Author contributions

Machine learning-aided segmentation F.B and F.Y; Cell Classification and synapse identification F.B.; CAM Process Caliber and Distance Estimation, Y.M., Figures, F.B; Statistical analyses., F.B. and Y.M.; 3D Renderings, F.B; Writing, F.B and B.H.; Review, Y.M.; Funding acquisition, J.W.L and B.H.; Guidance, J.W.L

## Acknowledgement

This work was supported by Human Frontier Science Program (HFSP) grant no. RGP0042/2019-102, Israel Science Foundation (ISF) grant no. 1928/15, National Institutes of Health (NIH) grant no. 5U24 NS109102 and NIH U01 NS108637. This work was also supported by a traveling fellowship from the Aharon and Ephraim Katzir Study Grants to F. Bidel.

## Bibliography

Babadi, B., Sompolinsky, H., 2014. Sparseness and Expansion in Sensory Representations. Neuron 83, 1213–1226. 10.1016/j.neuron.2014.07.035

Bartkowska, K., Tepper, B., Turlejski, K., Djavadian, R., 2022. Postnatal and Adult Neurogenesis in Mammals, Including Marsupials. Cells 11, 2735. 10.3390/cells11172735

Berger, D.R., Seung, H.S., Lichtman, J.W., 2018. VAST (Volume Annotation and Segmentation Tool): Efficient Manual and Semi-Automatic Labeling of Large 3D Image Stacks. Front Neural Circuits 12. 10.3389/fncir.2018.00088

Bidel, F., Meirovitch, Y., Schalek, R.L., Lu, X., Pavarino, E.C., Yang, F., Peleg, A., Wu, Y., Shomrat, T., Berger, D.R., Shaked, A., Lichtman, J.W., Hochner, B., 2023. Connectomics of the Octopus vulgaris vertical lobe provides insight into conserved and novel principles of a memory acquisition network. eLife 12, e84257. 10.7554/eLife.84257

Boycott, B.B., Young, J.Z., 1955. A Memory System in Octopus vulgaris Lamarck. Proceedings of the Royal Society of London B: Biological Sciences 143, 449–480. 10.1098/rspb.1955.0024

Budelmann, B.U., 1995. The cephalopod nervous system: What evolution has made of the molluscan design, in: Breidbach, P.D.D.O., Kutsch, P.D.W. (Eds.), The Nervous Systems of Invertebrates: An Evolutionary and Comparative Approach, Experientia Supplementum. Birkhäuser Basel, pp. 115– 138.

Bullock, TheodoreH., Budelmann, BerndU., 1991. Sensory evoked potentials in unanesthetized unrestrained cuttlefish: a new preparation for brain physiology in cephalopods. J Comp Physiol A 168, 141–150. 10.1007/BF00217112

Cervantes, D.C., Khare, H., Wilson, A.M., Mendoza, N.D., Coulon--Mahdi, O., Lichtman, J.W., Zurzolo, C., 2022. 3D reconstruction of the cerebellar germinal layer reveals intercytoplasmic connections between developing granule cells. 10.1101/2022.08.21.504684

Fiorito, G., Chichery, R., 1995. Lesions of the vertical lobe impair visual discrimination learning by observation in< i> Octopus vulgaris</i>. Neuroscience letters 192, 117–120.

Gould, E., Beylin, A., Tanapat, P., Reeves, A., Shors, T.J., 1999. Learning enhances adult neurogenesis in the hippocampal formation. Nat Neurosci 2, 260–265. 10.1038/6365

Hayworth, K.J., Morgan, J.L., Schalek, R., Berger, D.R., Hildebrand, D.G.C., Lichtman, J.W., 2014. Imaging ATUM ultrathin section libraries with WaferMapper: a multi-scale approach to EM reconstruction of neural circuits. Frontiers in Neural Circuits 8. 10.3389/fncir.2014.00068

Kier, W.M., 1996. Muscle development in squid: Ultrastructural differentiation of a specialized muscle fiber type. J Morphol 229, 271–288. 10.1002/(SICI)1097-4687(199609)229:3<271::AID-JMOR3>3.0.CO;2-1

Lin, A.C., Bygrave, A.M., de Calignon, A., Lee, T., Miesenböck, G., 2014. Sparse, decorrelated odor coding in the mushroom body enhances learned odor discrimination. Nature Neuroscience 17, 559–568. 10.1038/nn.3660

Litwin-Kumar, A., Harris, K.D., Axel, R., Sompolinsky, H., Abbott, L.F., 2017. Optimal Degrees of Synaptic Connectivity. Neuron 93, 1153-1164.e7. 10.1016/j.neuron.2017.01.030

Pavarino, E.C., Yang, E., Dhanyasi, N., Wang, M.D., Bidel, F., Lu, X., Yang, F., Francisco Park, C., Bangalore Renuka, M., Drescher, B., Samuel, A.D.T., Hochner, B., Katz, P.S., Zhen, M., Lichtman, J.W., Meirovitch, Y., 2023. mEMbrain: an interactive deep learning MATLAB tool for connectomic segmentation on commodity desktops. Frontiers in Neural Circuits 17.

Rochefort, C., Gheusi, G., Vincent, J.-D., Lledo, P.-M., 2002. Enriched odor exposure increases the number of newborn neurons in the adult olfactory bulb and improves odor memory. J Neurosci 22, 2679– 2689. 10.1523/JNEUROSCI.22-07-02679.2002

Sakamoto, M., Imayoshi, I., Ohtsuka, T., Yamaguchi, M., Mori, K., Kageyama, R., 2011. Continuous neurogenesis in the adult forebrain is required for innate olfactory responses. Proc Natl Acad Sci U S A 108, 8479–8484. 10.1073/pnas.1018782108

Sanders, G.D., 1975. The Cephalopods. In: Corning W.C., Dyal J.A. & Willows A.O.D. (eds) Invertebrate learning, Vol. 3. Plenium Press, New-York, pp 1–101.

Shani-Narkiss, H., Vinograd, A., Landau, I.D., Tasaka, G., Yayon, N., Terletsky, S., Groysman, M., Maor, I., Sompolinsky, H., Mizrahi, A., 2020. Young adult-born neurons improve odor coding by mitral cells. Nat Commun 11, 5867. 10.1038/s41467-020-19472-8

Shomrat, T., Turchetti-Maia, A.L., Stern-Mentch, N., Basil, J.A., Hochner, B., 2015. The vertical lobe of cephalopods: an attractive brain structure for understanding the evolution of advanced learning and memory systems. Journal of Comparative Physiology A 201, 947–956. 10.1007/s00359-015-1023-6

Shomrat, T., Zarrella, I., Fiorito, G., Hochner, B., 2008. The Octopus Vertical Lobe Modulates Short-Term Learning Rate and Uses LTP to Acquire Long-Term Memory. Current Biology 18, 337–342. 10.1016/j.cub.2008.01.056

Turchetti-Maia, A., Shomrat, T., Hochner, B., 2017. The Vertical Lobe of Cephalopods—A Brain Structure Ideal for Exploring the Mechanisms of Complex Forms of Learning and Memory. The Oxford Handbook of Invertebrate Neurobiology. 10.1093/oxfordhb/9780190456757.013.29

Turchetti-Maia, A.L., Stern-Mentch, N., Bidel, F., Nesher, N., Shomrat, T., Hochner, B., 2018. A novel NO-dependent ‘molecular-memory-switch’ mediates presynaptic expression and postsynaptic maintenance of LTP in the octopus brain. bioRxiv 491340. 10.1101/491340

Vicidomini, C., Guo, N., Sahay, A., 2020. Communication, Cross Talk, and Signal Integration in the Adult Hippocampal Neurogenic Niche. Neuron 105, 220–235. 10.1016/j.neuron.2019.11.029

Young, J.Z., 1963. The Number and Sizes of Nerve Cells in Octopus. Proceedings of the Zoological Society of London 140, 229–254. 10.1111/j.1469-7998.1963.tb01862.x

